# XCVATR: Characterization of Variant Impact on the Embeddings of Single -Cell and Bulk RNA-Sequencing Samples

**DOI:** 10.1101/2021.06.01.446668

**Authors:** Arif O Harmanci, Akdes Serin Harmanci, Tiemo Klisch, Akash J Patel

**Affiliations:** University of Texas Health Science Center, School of Biomedical Informatics, Center for Precision Health, Houston, TX 77030; Jan and Dan Duncan Neurological Research Institute, Texas Children’s Hospital, Houston, TX 77030; Department of Molecular and Human Genetics, Baylor College of Medicine, Houston, TX 77030; Department of Neurosurgery, Baylor College of Medicine, Houston, TX 77030

## Abstract

Gene expression profiling via RNA-sequencing has become standard for measuring and analyzing the gene activity in bulk and at single cell level. Increasing sample sizes and cell counts provides substantial information about transcriptional architecture of samples. In addition to quantification of expression at cellular level, RNA-seq can be used for detecting of variants, including single nucleotide variants and small insertions/deletions and also large variants such as copy number variants. The joint analysis of variants with transcriptional state of cells or samples can provide insight about impact of mutations. To provide a comprehensive method to jointly analyze the genetic variants and cellular states, we introduce XCVATR, a method that can identify variants, detect local enrichment of expressed variants, within embedding of samples and cells. The embeddings provide information about cellular states among cells by defining a cell-cell distance metric. Unlike clustering algorithms, which depend on a cell-cell distance and use it to define clusters that explain cell clusters globally, XCVATR detects the local enrichment of expressed variants in the embedding space such that embedding can be computed using any type of measurement or method, for example by PCA or tSNE of the expression levels. XCVATR searches local patterns of association of each variant with the positions of cells in an embedding of the cells. XCVATR also visualizes the local clumps of small and large-scale variant calls in single cell and bulk RNA-sequencing datasets. We perform simulations and demonstrate that XCVATR can identify the enrichments of expressed variants. We also apply XCVATR on single cell and bulk RNA-seq datasets and demonstrate its utility.

## Introduction

Gene expression profiling experiments generate large datasets that contain information about the activity levels of all genes in the transcriptome for large number of samples. The analysis of the complex and high dimensional data can help identify hidden patterns in the expression levels of driver genes, such as disease markers^1,2^, and delineation of transcriptional architecture of development. These results help formulate new hypotheses and perform validations^3^. RNA-sequencing is the standard approach for profiling gene expression in large samples^4^, whereby the cDNA from the isolated RNA is sequenced to provide estimates of expression levels of the genes. Unlike gene expression arrays, RNA sequencing provides much more information than estimation of expression levels such as allele-specific expression and eQTL mapping^5–7^, detection of small variants^8^, large copy number detection^9^, and transcriptional dynamics^10,11^. The decreasing cost of DNA-sequencing enables increasing the sample sizes and thousands of samples are profiled^12^ with hundreds of tissues^13,14^. In parallel to the samples sizes, there are now the single cell technologies^15–17^ whereby the expression can be measured at the single cell level for thousands of cells for hundreds of cell types and cellular states. In turn, the amount of information that needs be summarized and interpreted is increasing at a challenging pace.

One of the main challenges of analyzing these large datasets is efficient summarization of the biological information. To analyze thousands of samples or cells in a meaningful manner^18^, one of the first steps in the analysis is decreasing dimensionality by embedding the cells, such as PCA or tSNE. The embedding enables summarization of the transcriptomic state of the cells and put them in a simple perspective so that they can be clustered^19^, differential-expression can be assessed among clusters^20^, cell types can be assigned^21^, and they can be integrated with other datasets across multiple modalities^22^. While numerous embedding techniques are proposed^23^, these techniques are sometimes only used for visualization purposes only and there is much work that needs to be performed to utilize the embedding space in downstream analysis.

In this paper, we focus on integration and interpretation of the genetic variation within the embedding space that is used to analyze single cell and bulk RNA-sequencing datasets. The main motivation for developing a new method stems from our observations that cells that are mutated with similar mutations are generally cluster together in “clumps” that can provide interesting insight. For example, a driver mutation can manifest a specific transcriptional state on the cells that harbor the mutation. These cells, are then expected to form the clumps in the embedding coordinates. The most consequential variants are the large scale CNVs that clearly show clump patterns in tSNE and PCA embeddings of gene expression levels. XCVATR aims at systematically identifying these expressed variant clumps. Our approach, named XCVATR, is a flexible and integrated framework for detecting, filtering, and analyzing the association of the mutations (i.e. enrichment of mutations on embedding space) with the distances that are defined by the cell-embedding techniques. There are two main aspects that XCVATR is different from clustering methods that use variant calls to cluster cells^24,25^: First, XCVATR utilizes the an existing embedding and maps the variant allele frequencies on the embedding and detect local patterns of enrichment of the expressed alleles, i.e., spatial-correlation between the expressed variant^26^ alleles frequencies and embedding coordinates. This is different from the clustering methods that define the distance metric using the variants themselves. Secondly, XCVATR identifies local patterns, unlike the clustering algorithms that aim at finding a clustering of the cells that optimizes the global clustering of the cells. In addition, XCVATR sets out to be a self-contained framework for detection, annotation, and filtering of small variants, and detection and visualization the association of variant allele frequencies in the embedding space. This way, there is no dependency on other methods and the parameters of variant calling and filtering can be explicitly controlled.

One of the major components of XCVATR is the embedding that is used to summarize the transcriptomic states of the cells. The embedding is used to define the cell-cell distances. XCVATR expects the embedding to preserve locality information, i.e., the cells that are close to each other in the embedding coordinates are biologically similar to each other. This is a reasonable expectation for number of dimensionality reduction techniques such as PCA, tSNE, and UMAP. Among these, tSNE and UMAP probabilistically preserve locality, i.e., there is a random component in the embedding. We analyze and demonstrate evidence that the locality information is fairly well-preserved even in the presence of randomness in the embedding. It should also be noted that the opposite statement does not have to hold, i.e., we do not expect all cells that are biologically similar to map close to each other in the embedding space. This is a strict requirement that would require the embedding to preserve biological information almost exactly^27^ – it is, however, not necessary to analyze the association of the variants with embedding coordinates and geometry.

Overall, XCVATR combines various components of the general analysis steps into one flexible package for identifying, filtering, and visualizing the variants at different levels (reads, variants, embedding) and also analyzes the spatial distributions. This way, it represents a one-stop-shop for detecting. Single cell data makes it challenging to combine these steps.

## Results

We first overview the XCVATR algorithm then we present the main results.

### Overview of XCVATR Algorithm

Figure 1 summarizes the steps of XCVATR algorithm. The input to XCVATR is a mapped read file (such as a BAM file). While this is natural for single cell data, the bulk datasets contain many BAM files (one for each replicate), which can also be used in the analysis without and extra pre-processing. These steps are summarized here (See Methods for details).

**Figure 1.**
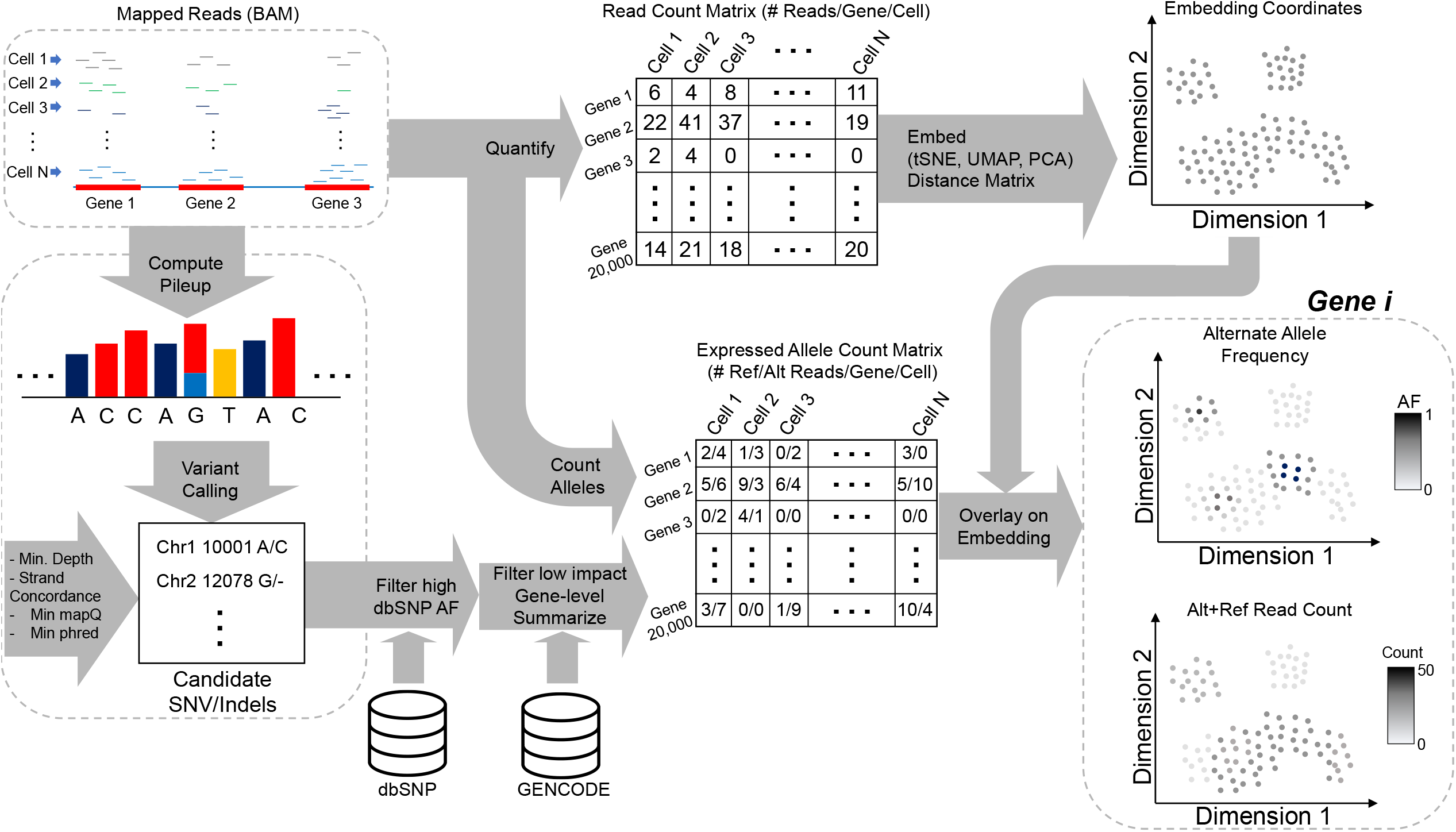
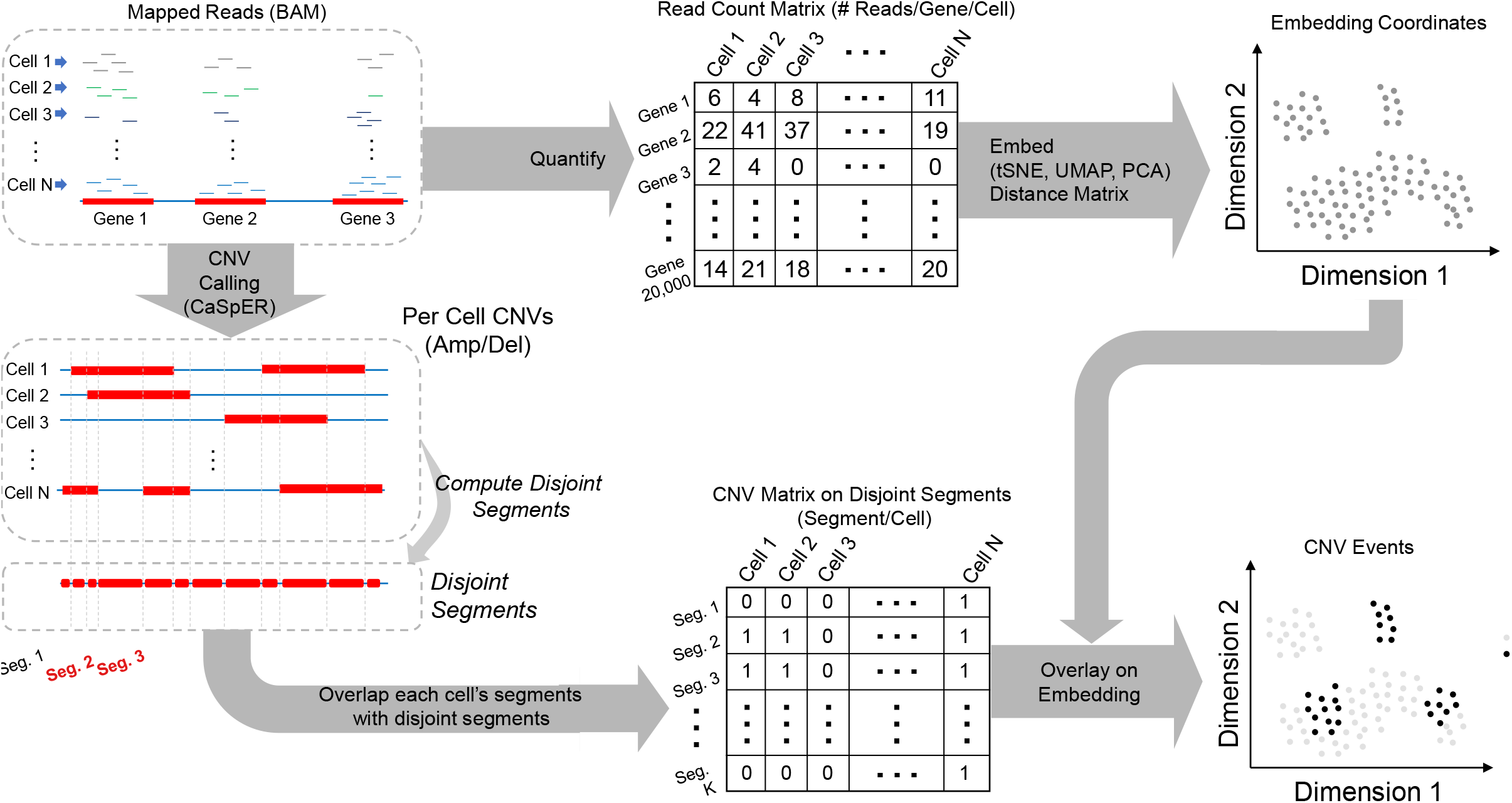
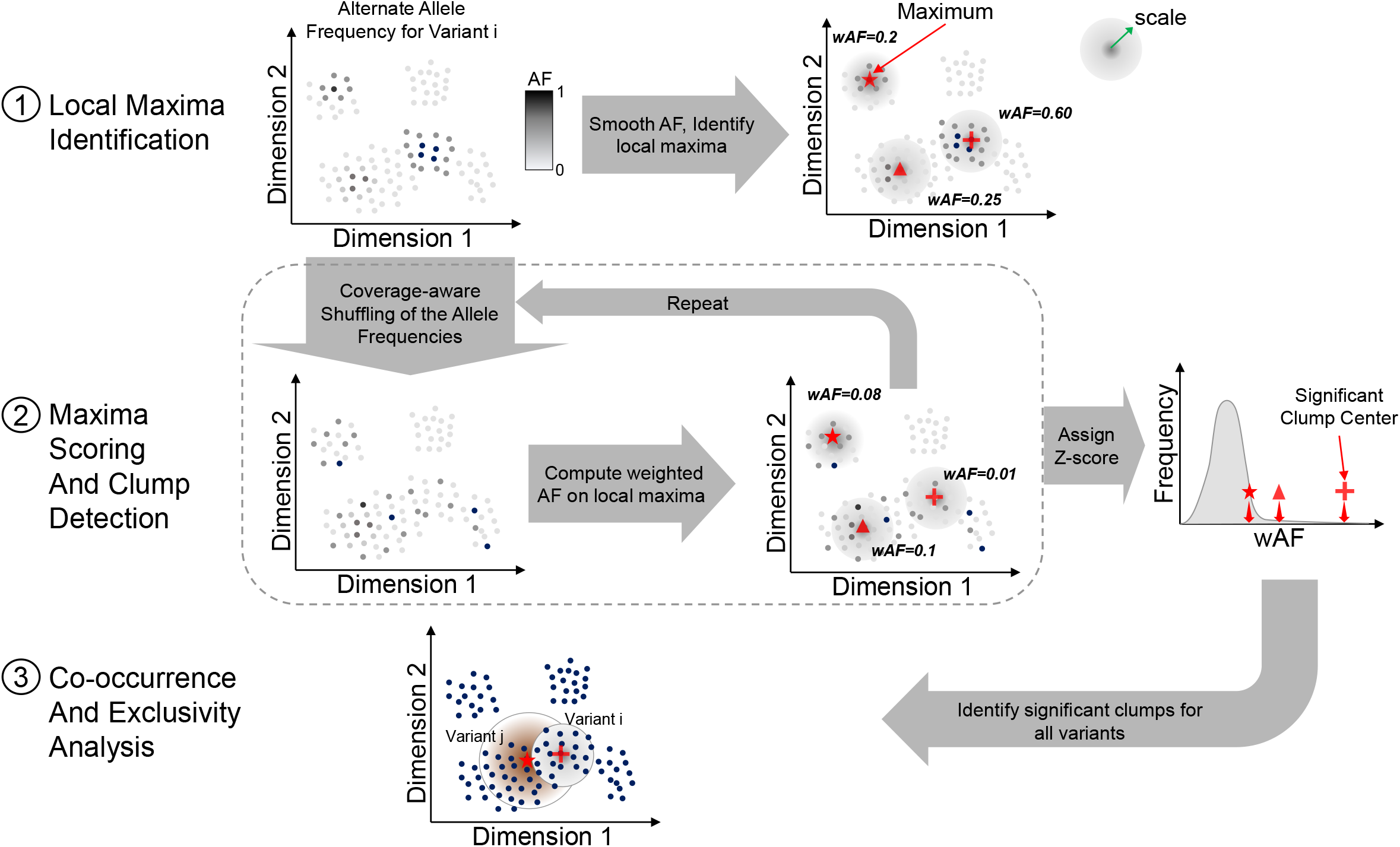
The steps of XCVATR algorithm.

XCVATR relies on the distance matrix between cells based on the transcriptomic profiles of cells (or samples). To generate the distance matrix, the first step is read count quantification for each cell (or samples), which are used for computing either the embedding coordinates of the cells or building the distance matrix directly from the expression levels.

Next, XCVATR performs detection and annotation of the genetic variants. The variant detection is designed to include an integrated and flexible SNV/indel calling step into XCVATR’s variant clump detection analysis. The variant detection can be parametrized in a relaxed manner so that the users can evaluate the variant clumps for variants that may be missed with conventional pipelines (i.e., variants with low allele frequencies, etc). We hypothesized that a relaxed variant calling can be meaningful since the variants will be further filtered in the context of variant clumping analysis. We therefore suggest the variant calls from XCVATR should not be used for other types of downstream analysis other than clumping analysis. If there is a variant call set (i.e. VCF file) generated by other pipelines (GATK^28^, Mutect^29^), the users can provide these as input and skip the variant detection step. XCVATR uses pileups to identify candidates SNVs that are passed through several filters. This is a strategy similar to that used by VarScan suite of variant callers^30,31^.

Variant annotation is integrated into XCVATR to make the workflows more flexible and complete. Although there are well-established protocols and for variant annotation such as VEP^32^ and AnnoVar^33^, these methods occasionally change over time and it becomes challenging to integrate the output of these tools and provide reproducibility. To get around these, XCVATR performs variant annotation step to provide a flexible filter to select variants with respect to impact. XCVATR takes variant annotation file (GTF or GFF) and annotates the variants with respect to their impact on the protein sequence. These variants are then annotated and filtered to use the most impactful mutations (See Methods for details of annotation and variant calling on filters).

#### Allele Counting

For SNVs and small indels, XCVATR counts the number of mapping reads, for each read, that support the alternate and reference alleles. This is used to generate the estimated alternative allele frequency of each variant. XCVATR treats the alternate allele frequency of a variant These are then used as scores for a variant’s existence on each cell.

For copy number variants (CNVs), the variants are first separated into amplifications and deletions (Fig 1c). CNVs are different from small variants since they can cover large domains as long as the chromosomal arms. To analyze different length scales, XCVATR performs clumping analysis for large scale (chromosome arm length) and also at segment level scale. The large scale CNV analysis, there are 44 possible events for each deletion and amplification. For these, XCVATR first builds a binary count matrix that is analogous to the alternate allele frequency for the small variants. Each entry in this matrix indicates the existence of the CNV (row) in the corresponding cell (column). At the segment scale, each CNV is treated as a separate variant. However, since the CNVs identified in each cell has different coordinates, XCVATR first identifies the common amplification/deletion events by overlapping the CNVs from cells and identifying the minimal set of common variants (Fig 1). Next, these common variants are used to build a binary count matrix similar to the large-scale matrix. XCVATR analysis each of the common and disjoint events as a separate variant and performs variant clump analysis.

After the alleles are counted, XCVATR can optionally summarize the variants on genes by selecting the most impactful variant in each cell (or sample) om each protein coding gene. This can filter out and many variants and provide a clearer view on a gene-level to the user.

#### Smoothing Scale Selection on the Embedding

XCVATR performs a multi-scale analysis of the distances to identify the variant clumps. This resembles the multi-scale filters that are used to identify blobs in images^34–36^. Each scale defines a neighborhood around a cell in the embedding coordinates and is used to smooth the allele frequencies using a Gaussian filter that is centered on a cell and decreases as the cells get further from the center cell. The scales, however, must be tuned to the distance metric or the embedding coordinates. XCVATR performs a scale selection to tune the analysis to the selected cell-cell distance metric.

For each cell, XCVATR identifies *N*_*v*_ cells that are closest to the corresponding cell (i.e. neighbors). This defines the close neighborhood of each cell. XCVATR then scans the neighborhood size 

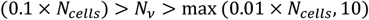

between 1% of the cells (or 10 cells, if lower) and 10% of the cells in the sample (*N*_*cells*_) and computes the radius of the neighborhood of each. For *N*_*v*_ cell neighborhood of a center cell, neighborhood radius is defined as the distance of the furthest cell to the current center cell. The minimum and maximum scales (*σ*_*min*_, *σ*_*max*_) are defined as the median of the neighborhood radii of all cells computed at the minimum and maximum neighborhood sizes *N*_*v*_ defined above (See Methods). The scales are used for smoothing the allele frequencies and identifying the variant clumps.

This computation can be performed efficiently since the distance matrix (unless it is provided) can be computed quickly from the embedding coordinates using fast matrix multiplications. Neighbor detection is performed by sorting the distances and selecting the closest *N*_*v*_ cell (or samples). After this step, only closest neighbors are processed by XCVATR.

#### Variant Clump Candidate Selection

One of the challenges in clump detection is the large number of cells that needs to be analyzed in different scales. To decrease the search space, XCVATR performs a cell-centered analysis, where XCVATR does not aim at modeling the whole embedding space but rather focuses on the cells, i.e., each detected clump is centered around a specific cell (or sample). We believe this is a reasonable expectation since the expected clumps are substantially larger than the cell-cell distances and therefore clump-detection should be accurate even when they are centered around cells. This way, XCVATR cuts the cost of modeling and searching the whole embedding space and focuses on cells.

Secondly, in visual evaluation of the variant allele frequency distributions on embeddings, the number of clumps were observed to be much smaller than the number of samples or cells. Motivated by this, we designed a candidate pre-selection that decreases the search space for the variant clumps. Given a smoothing scale *σ*_*a*_ at the scale *a*, XCVATR computes a smoothed AF value for each cell: 

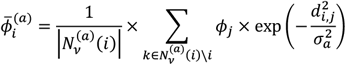

where *ϕ*_*j*_ denotes that alternate AF of the variant on the *j*^*th*^ cell (1> *ϕ*_*j*_ >0), *d*_*i,j*_ denotes the distance between *i*^*th*^ and *j*^*th*^ cells in the sample, 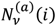 indicates the set of indices for the cells that are in the vicinity of the *i*^*th*^ cell for the scale *a*. From the above equation, the smoothed allele frequency of *i*^*th*^ cell, 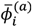, is higher when its neighborhood contains many cells with high allele frequencies. In addition, the smoothed AF depends on the scaling parameter *σ*_*a*_. Each scale is processed independently from other scales. The cells with high smoothed allele frequencies represent the potential variant clump centers in the embedding coordinates. XCVATR identifies the set of cells as candidates for which the cells in the neighborhood are strictly lower in terms of smoothed allele frequency (Figure 1): 

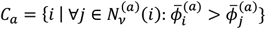

where *C*_*a*_ denotes the indices of cells that are clump centers. Above condition defines the candidate clump centers as the cells which have locally maximum smoothed allele frequencies when they are compared to the smoothed allele frequencies assigned to all their close neighbors.

#### Specification of Position and Size of Clumps

Up to current point, we identified a clump by the cell at its center, which specifies the position of the clump in the embedding space. In addition to the center, it is also necessary to define the radius of the clump so that the size of the clump can be specified. XCVATR makes use of the scale parameter at which the clump is identified, i.e., *σ*_*a*_ at scale *a*. Thus, all the cells that are closer than *σ*_*a*_ to the center of a clump are assigned to this clump. Later on, we provide a method to detect the most enriched

#### Variant Clump Evaluation by RD-aware permutation

For each of the cells in *C*_*a*_(*a*^*th*^ scale), the smoothed allele frequencies are compared to an empirical background. XCVATR utilizes a permutation test to assign significance to the each of the candidate clump centers. For this, XCVATR generates *K* permutations of the AF’s that are assigned to the cells and computes the smoothed AF for all the candidate. For each permutation, the smoothed AFs are computed for each candidate clump center. XCVATR then computes a z-score that is used to rank the clumps. 

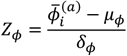

One of the important factors is to ensure that the AF provides new information and does not simply recapitulate the coordinates-based information in the embedding. One example for this is that some cells may exhibit a cell-specific marker that is not expressed in other cells at all. In this case, a germline variant will be expressed on these cells while other cells will have no expression of the variant. In such a scenario, a naïve approach would determine that this variant exhibits a clump on the cells where the gene is expressed. This would be an uninteresting clump that emerges based on the cell-type specificity of the gene. XCVATR aims at uncovering variant-specific clumps. To filter out these clumps, XCVATR sets a threshold *τ* on the total read depth at which the variant is expressed. XCVATR also reports the read depth z-score. This way, the clumps are evaluated with respect to the read-depth bias.

#### Read-level and Cell-level Enrichment of the Alternate Allele Expression in the Clumps

In order to filter the clumps, XCVATR computes the significance of enrichment of expressed alternate alleles at the read level and at the cell (or sample) level in each clump. To compute the enrichment at the read level, XCVATR first computes the total number of alternate and reference reads in all cells. These are used to estimate a baseline (bulk) alternate AF. Next, for each clump, the total alternate allele supporting reads and total reads are computed. At scale *a*, and the *b*^*th*^ clump, these are used to compute the read-level modified binomial p-value: 

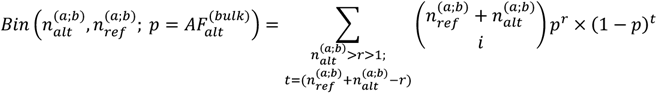

where 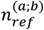 and 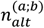 denote the number of reads that support alternate and reference alleles for the corresponding variant in the *b* clump that is identified in scale *a*: 

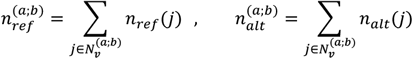

where 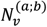 denotes the neighborhood of the center of the clump *b* at the scale *a*, and *n*_*ref*_(*j*) indicates the number of reference alleles in cell *j*. In above equation, the flip probability is chosen as 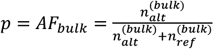, which represents the alternate allele frequency of the variant in the whole bulk sample. The binomial p-value estimates the enrichment of the alternate allele supporting reads in the clump *b* when compared to randomly assigning reads to all cells with probability *p* = *AF*_*bulk*_.

Next, XCVATR computes enrichment of alternate AF at cell level. At the scale *a*, XCVATR counts the cells in the clump *b* whose alternate allele frequencies are above *η*. Next, XCVATR counts the number of cells in the whole sample for which the alternate allele frequency is above *η*. These values are used to compute a significance of the enrichment of alternate alleles at cell level using Fisher’s exact test: 

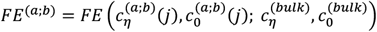

where 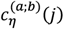 indicates the number of cells in clump *b* in scale *a* where allele frequency exceeds *η*: 

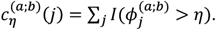

Similarly, 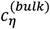 indicates the number of cells among all the cells (i.e., bulk) for which the allele frequency exceeds *η*. The read-level and cell-level enrichment estimates are used to filter out clumps that exhibit low levels of enrichment in comparison to the bulk sample at read and cell (or sample) level.

Finally, XCVATR computes the effective radius for each clump. For each clump, XCVATR iterates over the cells closest to the clump’s center cell. This way, the neighborhood around the clump’s center are analyzed as they expand in radius. For each neighborhood, the neighborhood where the cell-level enrichment is maximized (Fisher’s exact test p-value is minimized) is selected as the effective radius of the clump. After the clumps are identified, the clump centers and the scale at which they are identified, permutation z-scores, and alternate allele enrichment statistics, and effective radii are reported in the output.

#### Visualization

XCVATR provides visualization of the clumps on the embedding coordinates for each variant. This enables the users to manually evaluate the variants. This can also be helpful to visualize the cell-type specifications and phenotypic properties in comparison to the clumps. The visualization utilities are implemented in R and directly make use of the data generated by XCVATR.

### Analysis of Detected Variants and Testing of Clumps

To explore the statistics of the identified variants, we analyzed the alternate allele frequency of the detected variants in bulk (160 Meningioma patients^37^) and single cell datasets (BT-S2 sample from Darmanis et al.^38^). We observed generally that the detected variants exhibit allele frequency spectrum that are dominated by alternate AFs of 0% and 100% (Fig. 2) and a slight enrichment at 50%. For the single cell dataset, substantial portion of the mutations are expressed in small fraction of cells. This is in concordance with the previous studies^24^. In the single cell data, we also observed that most of the mutations are observed in small fraction of cells (Fig 2).

**Fig. 2:**
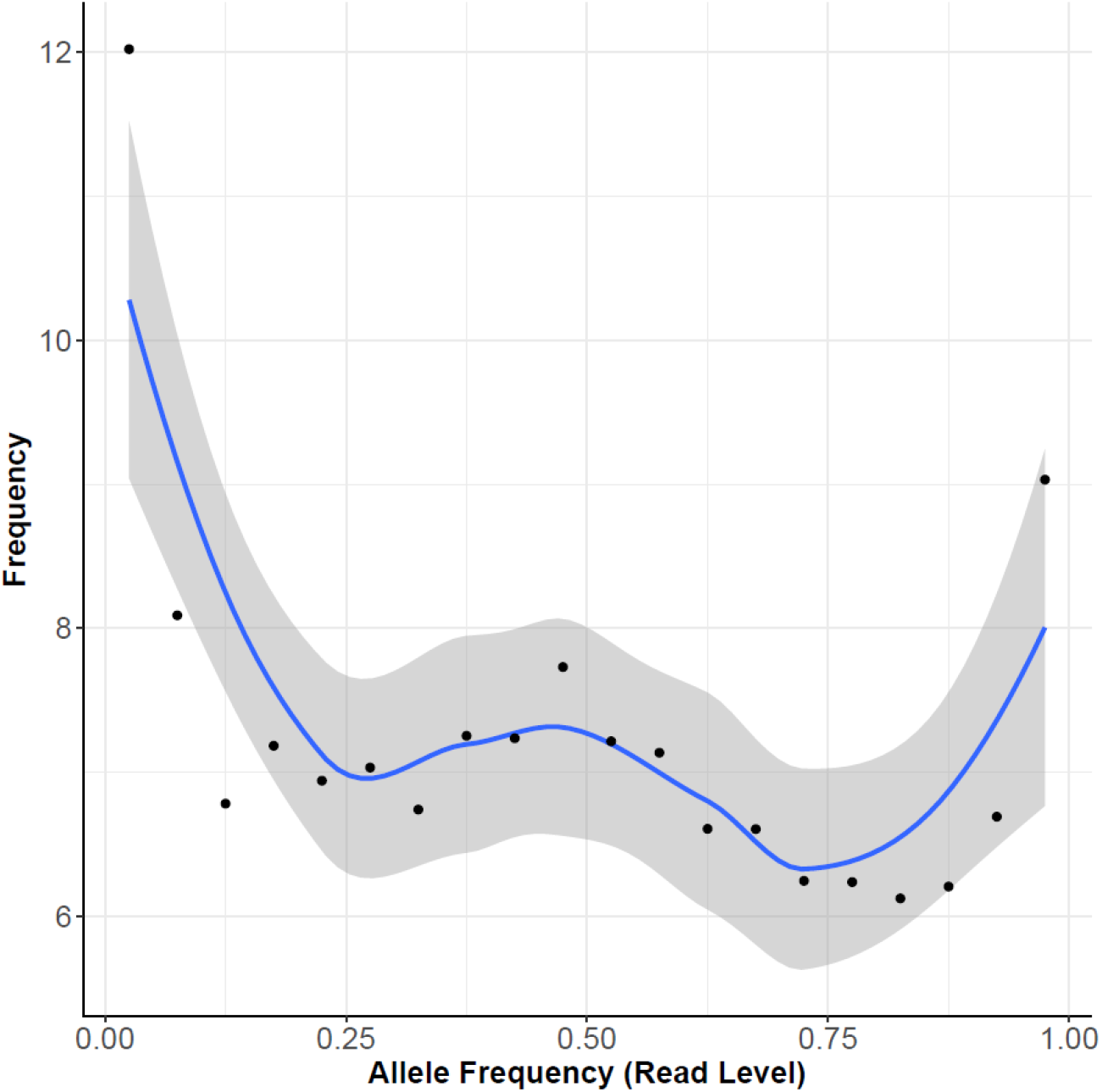
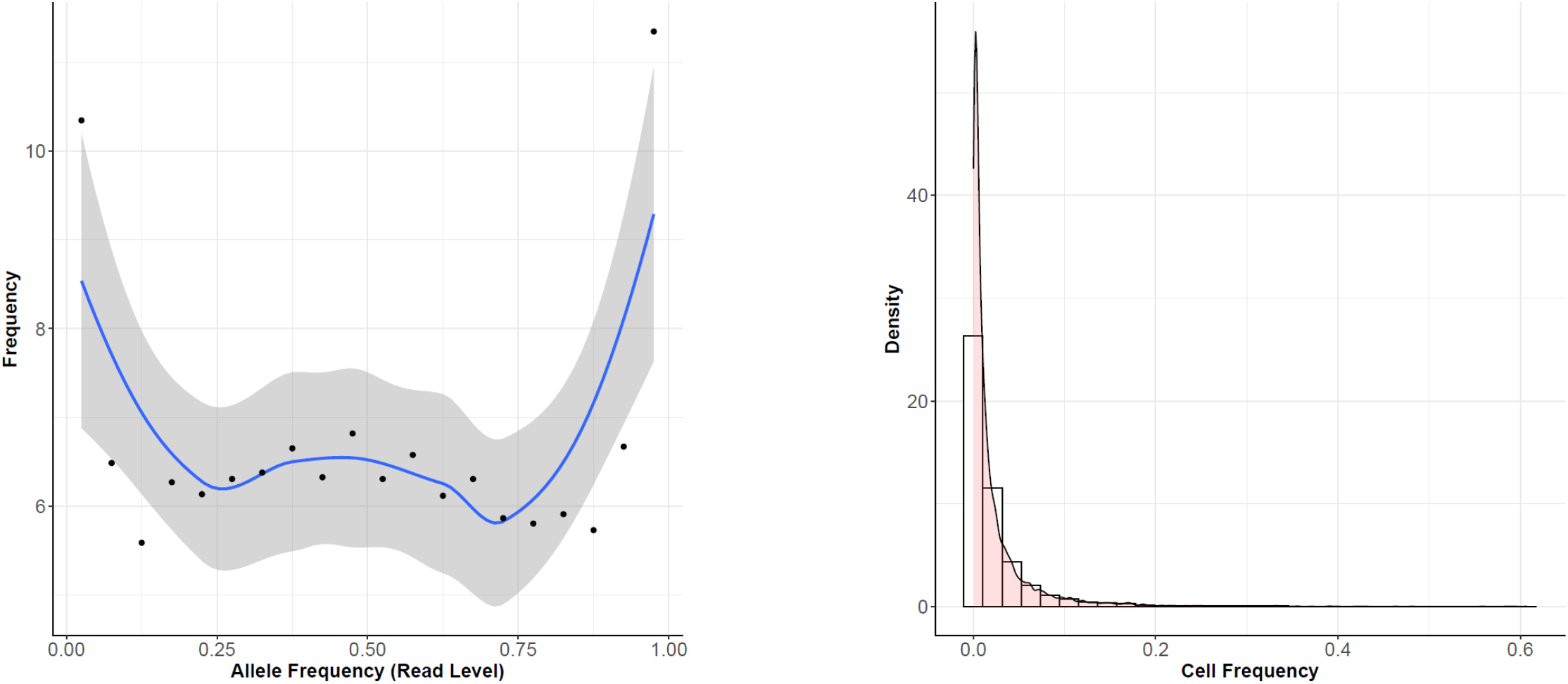
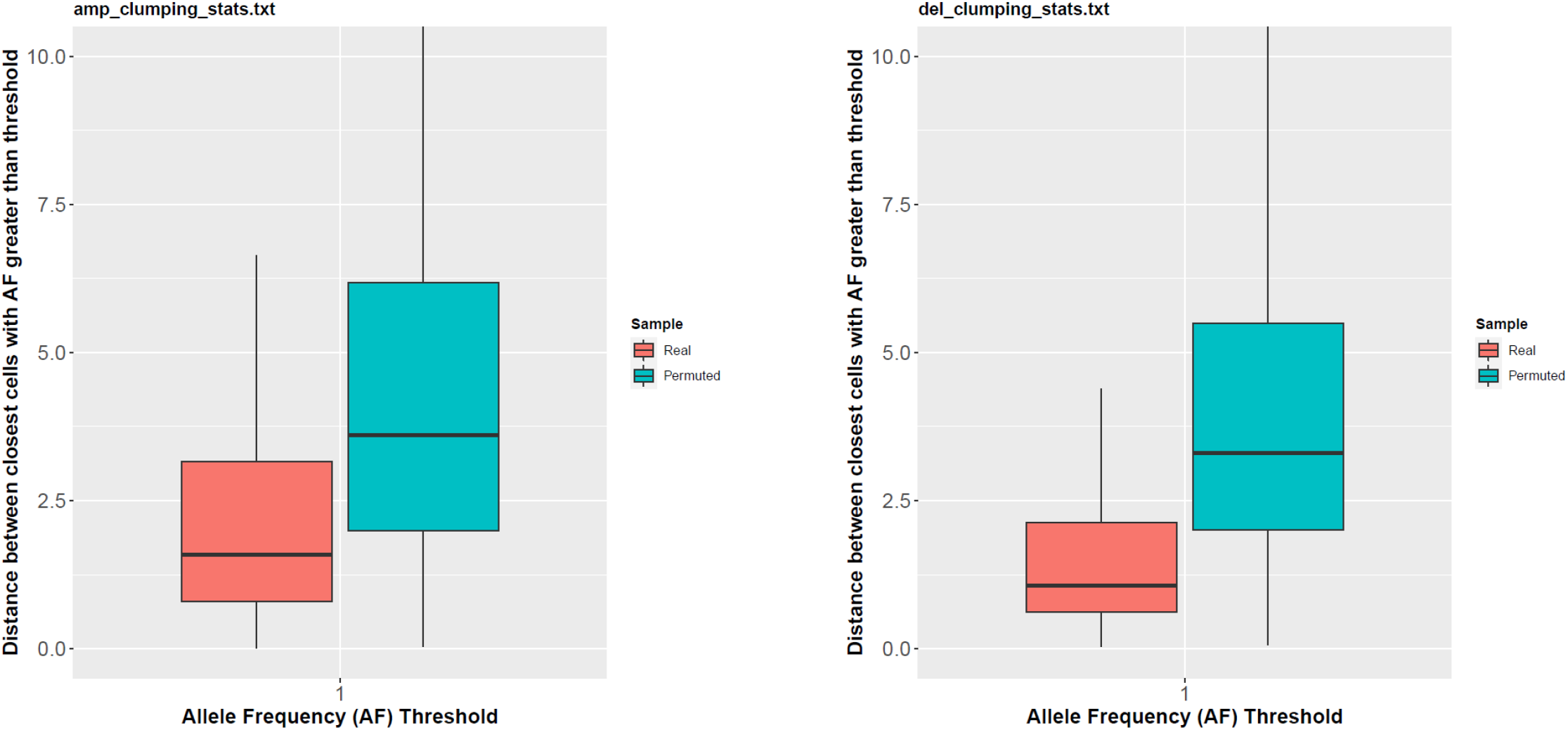
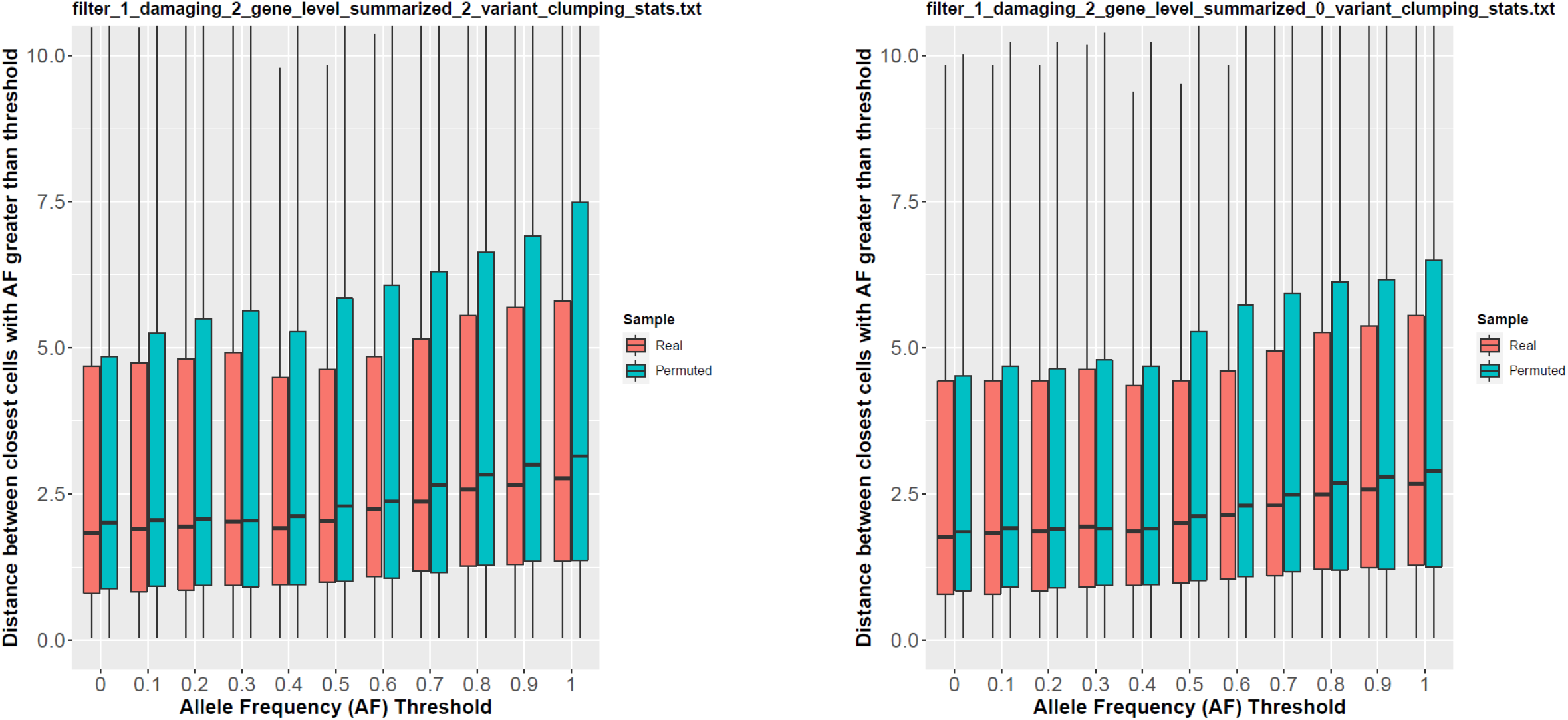
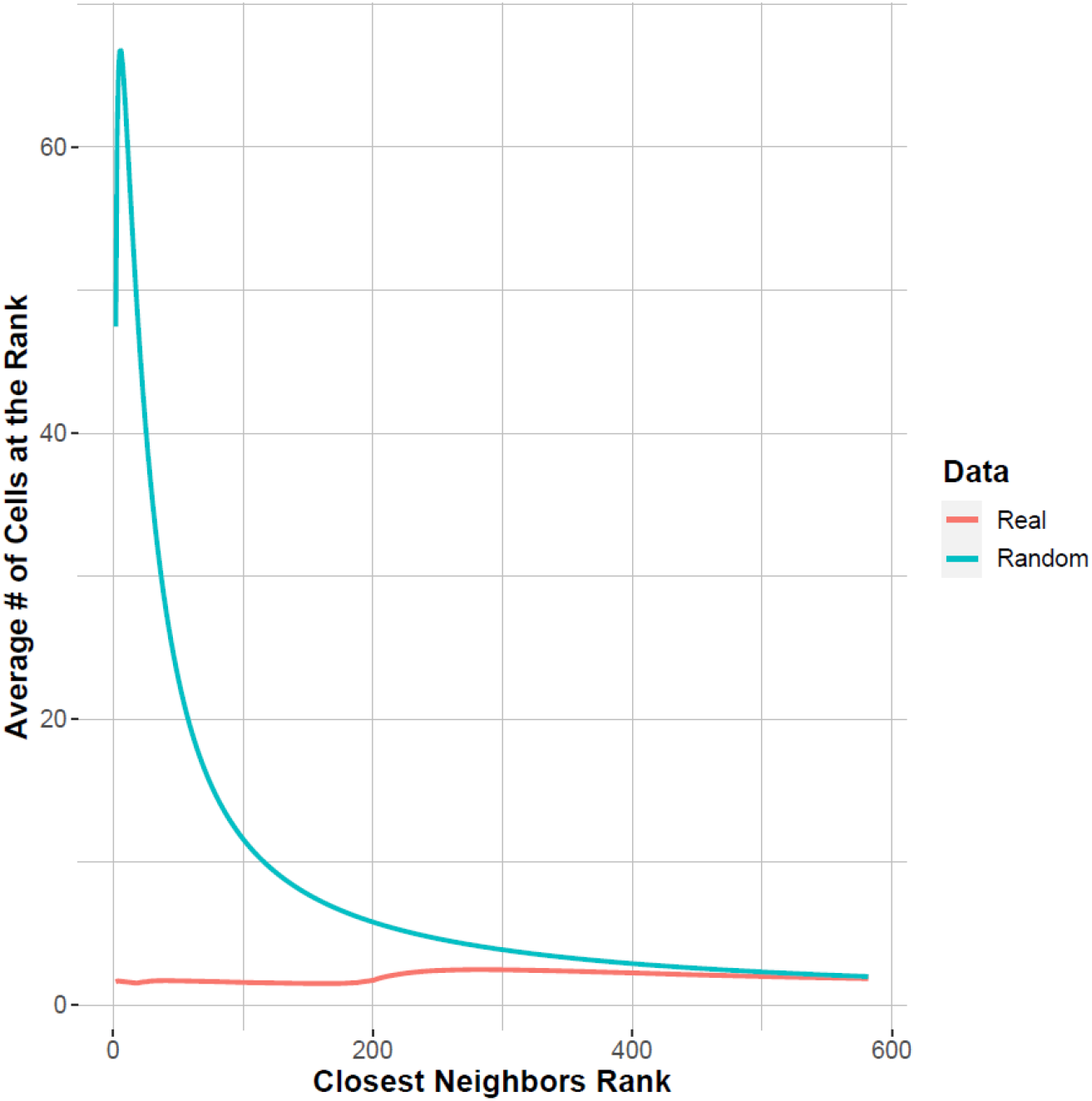
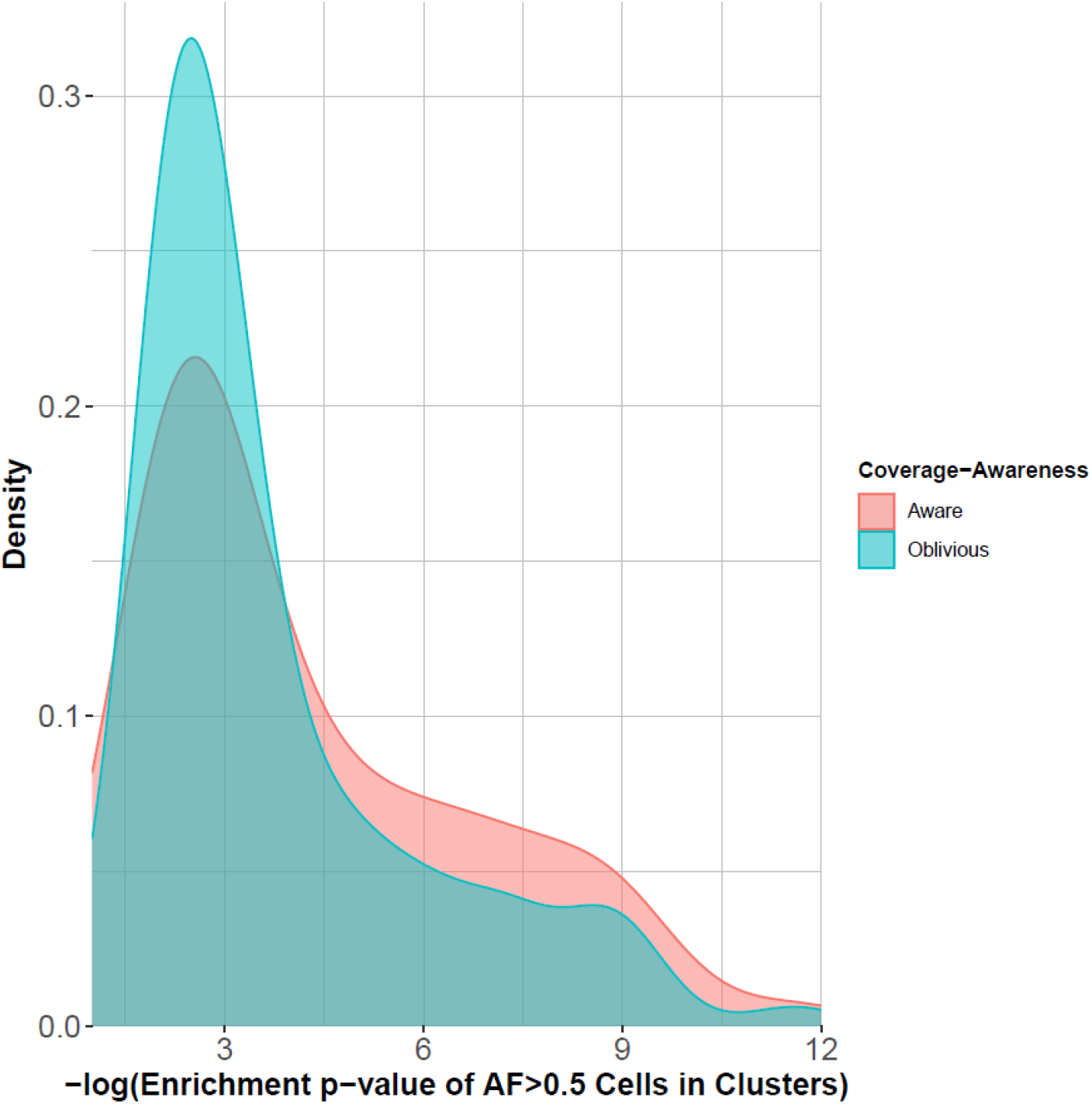
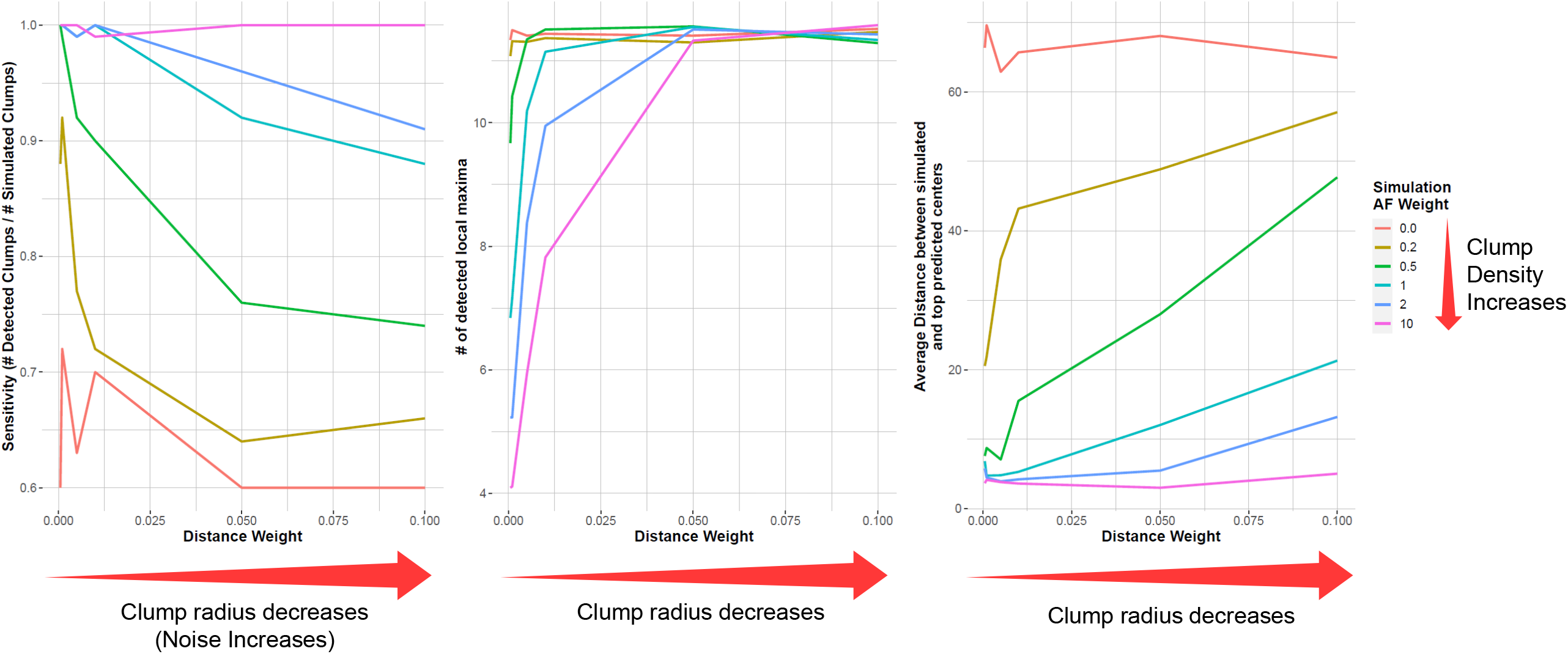
The analysis of clumping behavior and statistics of CNVs, SNVs. The conservation of local statistics by tSNE with different parameters and pseudorandom seeds.

We next quantified our hypothesis that variants clumps are observed in tSNE embeddings. Although we did manually observe clumps, it is useful to evaluate their existence automatically and objectively. To test whether the variant clumps frequently occur in embedding coordinates, we computed the average distance between cells that contain variants with high alternate allele frequencies. If these cells are closer to each of other than expected by random chance, this provides us with empirical evidence that there is detectable general clumping behavior of variants. We implemented the above procedure using Darmanis et al dataset. We computed the distribution of closest distance of cells that contain variants with alternate AF greater than *η*. We computed the same distribution in real and in shuffled data (Fig 2). As the allele frequencies cutoff, *η*, increases, we observed that the cells that contain variants with high alternate AFs are closer to each other compared to the shuffled data, where the allele frequencies are shuffled in read-depth aware manner. The clumping behavior is much clearer when we performed the same analysis with CNVs (Fig 2XX). For this, we computed the cell-cell distance distributions for the cells that contain CNVs (amplifications and deletions). Fig 2XX shows that the distribution of distances are much smaller compared to the shuffled datasets for both deletions and amplifications. The CNVs exhibit a much stronger clumping effect compared to the SNVs and indels since they potentially have much stronger impact on the transcriptional state of the cells. These results provide evidence for the potential general clumping of cells with respect to the SNV, indel, and CNV frequencies and they objectively motivate the need for a method such as XCVATR.

### Robustness of Local Statistics and Accuracy of Clump Detection

We evaluated the robustness of local distance statistics from the tSNE coordinates that we use as the embedding of the single cell RNA-seq data. This is generally necessary because tSNE has a random component that requires a seed pseudorandom-number generator^39^. When the seed is changed, or when tSNE is ran twice with the same seed, the returned coordinates completely change. We analyzed whether the closest samples are still close when the seed changes. The robustness of the locality of samples is essential for XCVATR’s clumping analysis since XCVATR aims at detecting the local clumping of the cells that harbor variants with high alternate AFs. To test the local robustness of the embedding coordinates, we first ran tSNE as provided by the SEURAT package^40^ with default parameters using the Darmanis et al. dataset. tSNE coordiantes are generated 100 times while changing the seed number with every run. Next, for each cell, we identified the neighborhoods upto 600 closest neighboring cells (out of 1,170 cells in total). For each of the 100 embeddings, we computed the number of unique cells at each of the 600 distinct closest neighborhoods and plotted the average number unique cells at each neighborhood size, which is shown in Figure 2. Compared to the shuffled dataset where we start with the shuffling of the cell identifiers coordinates, the real data shows strong preservation of the embedding. At each neighborhood size, which determines the rank or strength of locality, the real dataset contains on average between 1 and 3 unique cells for each closest cell. In addition, this average does not change with increasing neighborhood size. For shuffled data control data, the average number of unique cells for each closest cell increases to as high as 60 cells for each additional closest neighbor. It should be noted that these results only attest to the robustness of locality statistics. Over the neighborhoods covering more than 25% of the cells, the randomized and real locality statistics are very similar, which suggests that the clumps that contain larger than 25% of the cells may be non-robust and non-reproducible between different runs of tSNE.

Next, we tested the accuracy of multiscale clump detection approach. In order to have a ground truth, we simulated one clump and ran XCVATR to detect this simulated clump. For this, we simulated variant clumps on the tSNE embedding coordinates (See Methods) from the Darmanis et al. study. First, a cell is randomly selected and is designated as the known clump center. We simulated two parameters in the simulations. First is the scale parameter, which determines the size of the simulated clump. Second parameter is the strength of the clump (we refer to as “AF weight”), this is tuned by a parameter that enforces high AFs to be assigned closer to the center of the simulated clump. This parameter tunes how strong the AF is distributed around the clump. We simulated 5 different scales and 6 different AF weight parameters and for each parameter combination, we simulated 100 randomly selected clump centers. For each simulation, an alternate read count matrix is generated for the 1,170 cells in the simulated dataset and is input to XCVATR. The accuracy of the simulation is evaluated by comparison of the known clump center and the centers of the clumps detected by XCVATR. Any clump that is detected within less than 1% of the whole embedding space radius is deemed a match. We evaluated the fraction of times XCVATR was able to identify the cell at the center of the clumps correctly. We also recorded the number of clumps that are identified by XCVATR. Figure 2 shows these accuracy statistics.

We also evaluated how the RD-aware permutation impacts the identified clumps. For this, we identified the clumps from Darmanis et al. dataset with and without the RD-aware shuffling. We next plotted the distribution of the cell-level enrichment Fisher’s Exact test p-value from of the clumps detected with and without RD-aware shuffling (Fig. 2). We observed that the RD-aware shuffling enables detection of clumps with higher enrichment of cells with variants that are expressing higher alternate alleles. This analysis provides evidence that RD-aware shuffling can be helpful to identify clumps that are enriched more in cells expressed alternate alleles.

### Single Cell Glioma Datasets

For demonstration of data analysis using XCVATR, we first focused on the analysis of Darmanis et al, which contains single-cell RNA sequencing of 4 patients with glioma brain tumors using the Smart-Seq2 technology^41^. The advantage of Smart-Seq2 is that it provides a more uniform coverage compared to technologies such as Drop-Seq and 10X where there is a 3’-bias on the transcript^42^. While the bias can potentially bias the variant detection step, XCVATR does not require the variants to be complete or unbiased since each variant is analyzed independent of the other variants. The datasets were downloaded from GEO (Accession number GSE84465). We found 3590 cells from the metadata that were processed and mapped using Hisat2. We used the tSNE that is provided by the original study. Figure 3 shows the distribution of the cells from all the samples. After initial inspection of the samples, we focused on the sample with id “BT_S2” that contains the most impactful events in terms of the copy number variants, SNVs, and the indels. Also, this sample had 1170 cells and sequenced as one of the largest sample sequenced. Figure 3a shows an example of a clump that is identified by XCVATR for the deletion of chromosome arm 17q, which is identified as the top clump among other large-scale deletions. This is a well-known deletion that is observed in glioma tumors. Two clumps are highlighted on the malignant cells. Fig 3b highlights the deletion clumps on the 17p, 10q, and 22q arms, that are reported as the top CNVs. While these results are partially expected, these corroborate the clump detection performed by XCVATR.

**Fig. 3.**
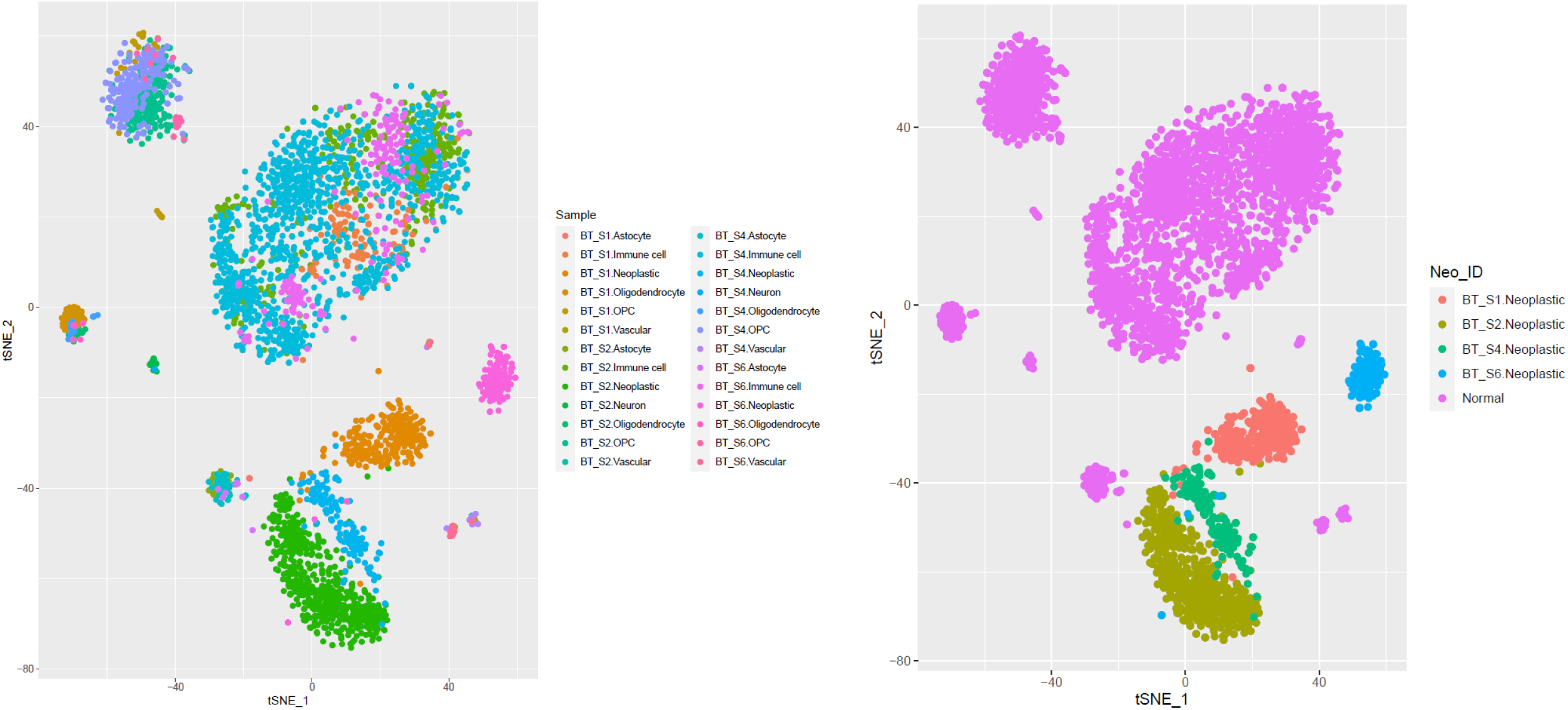
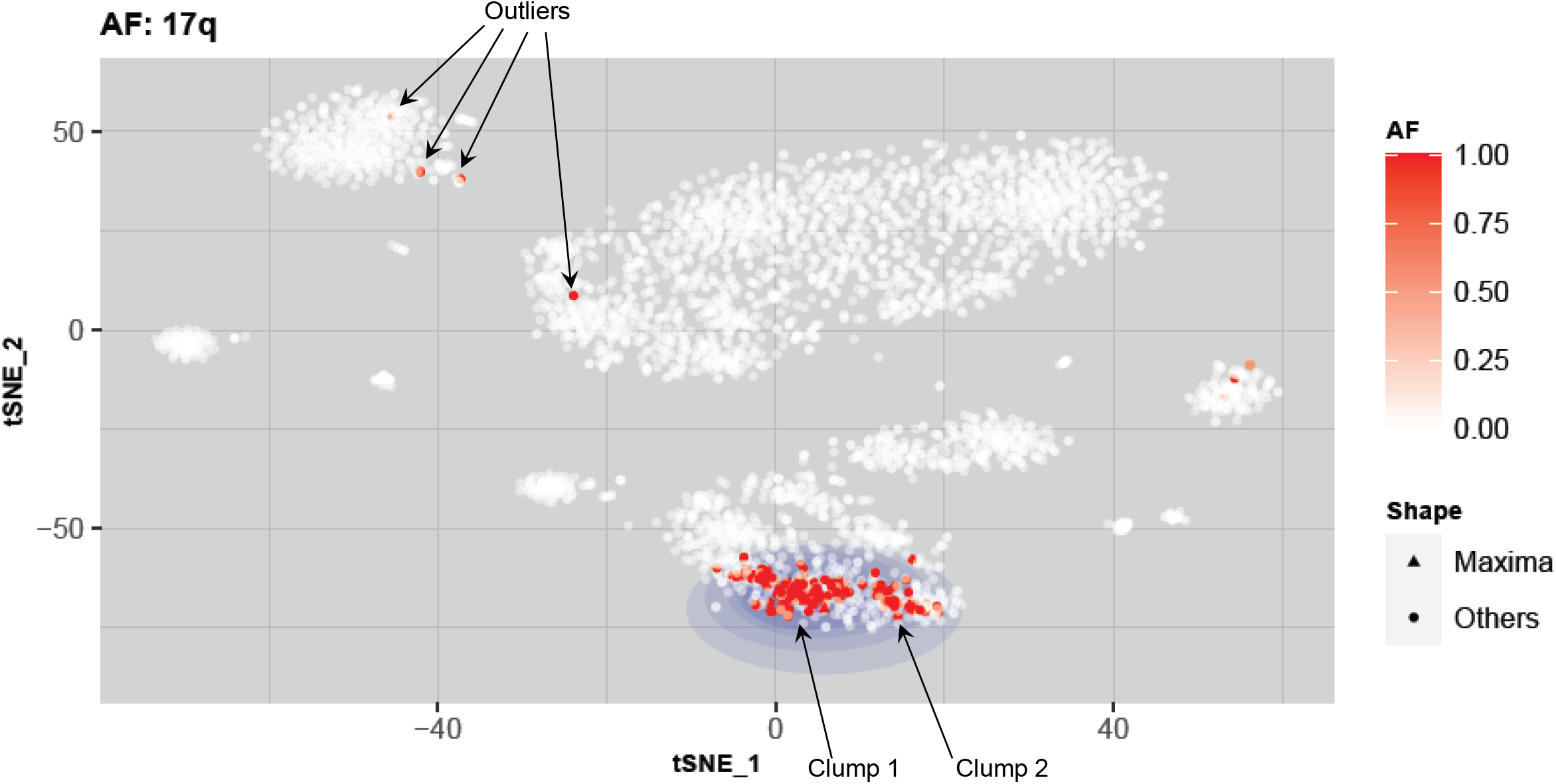
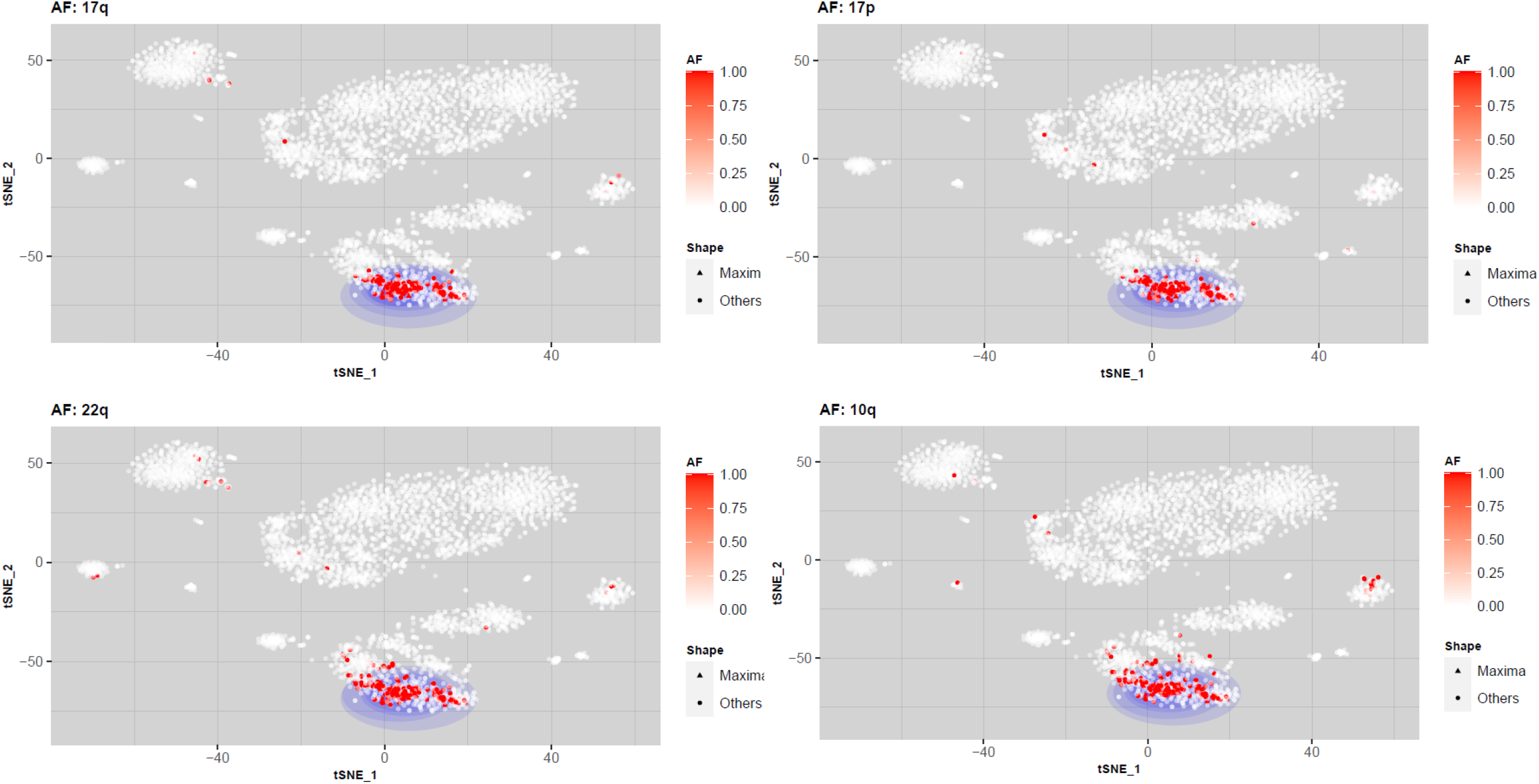
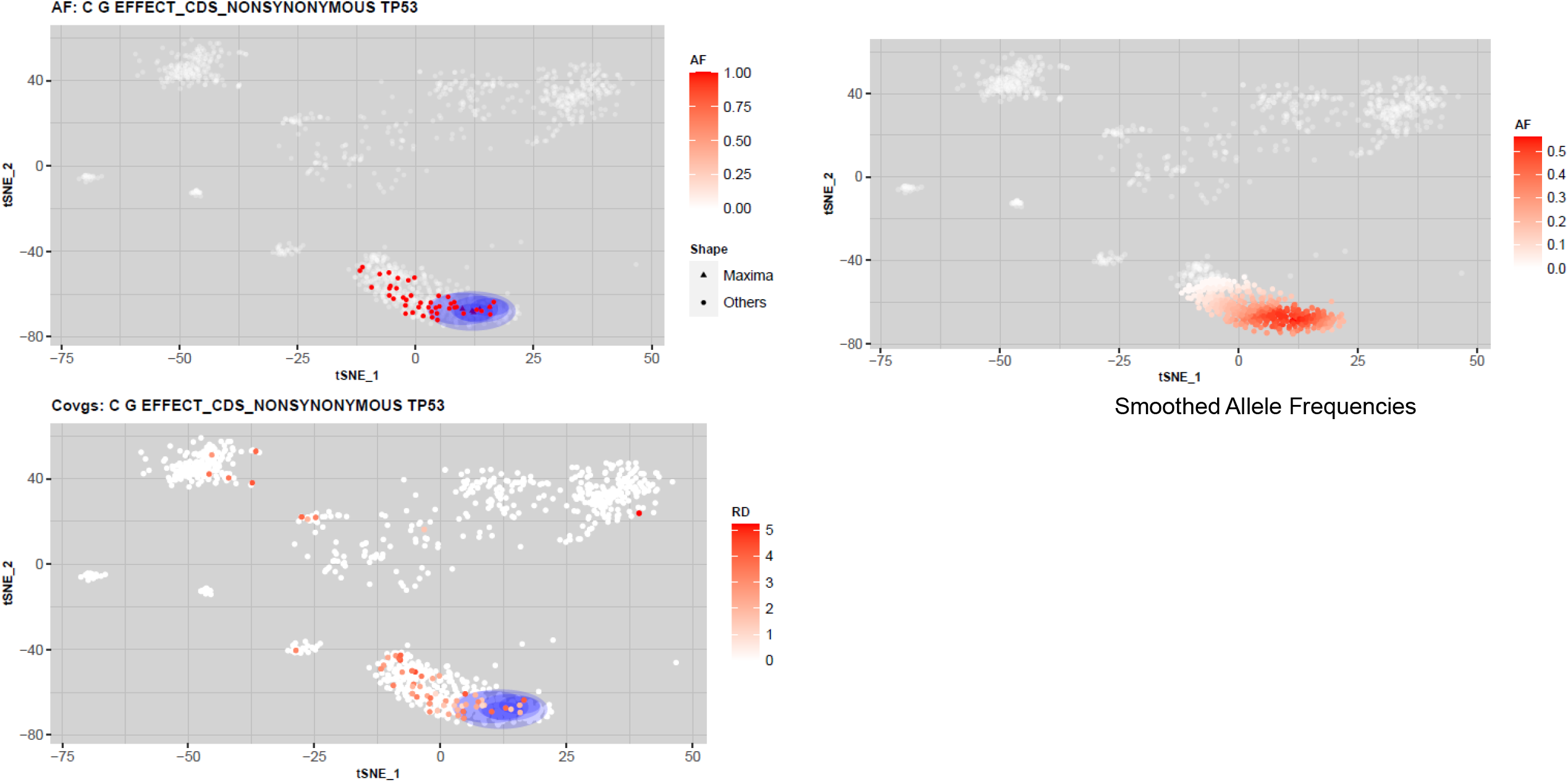
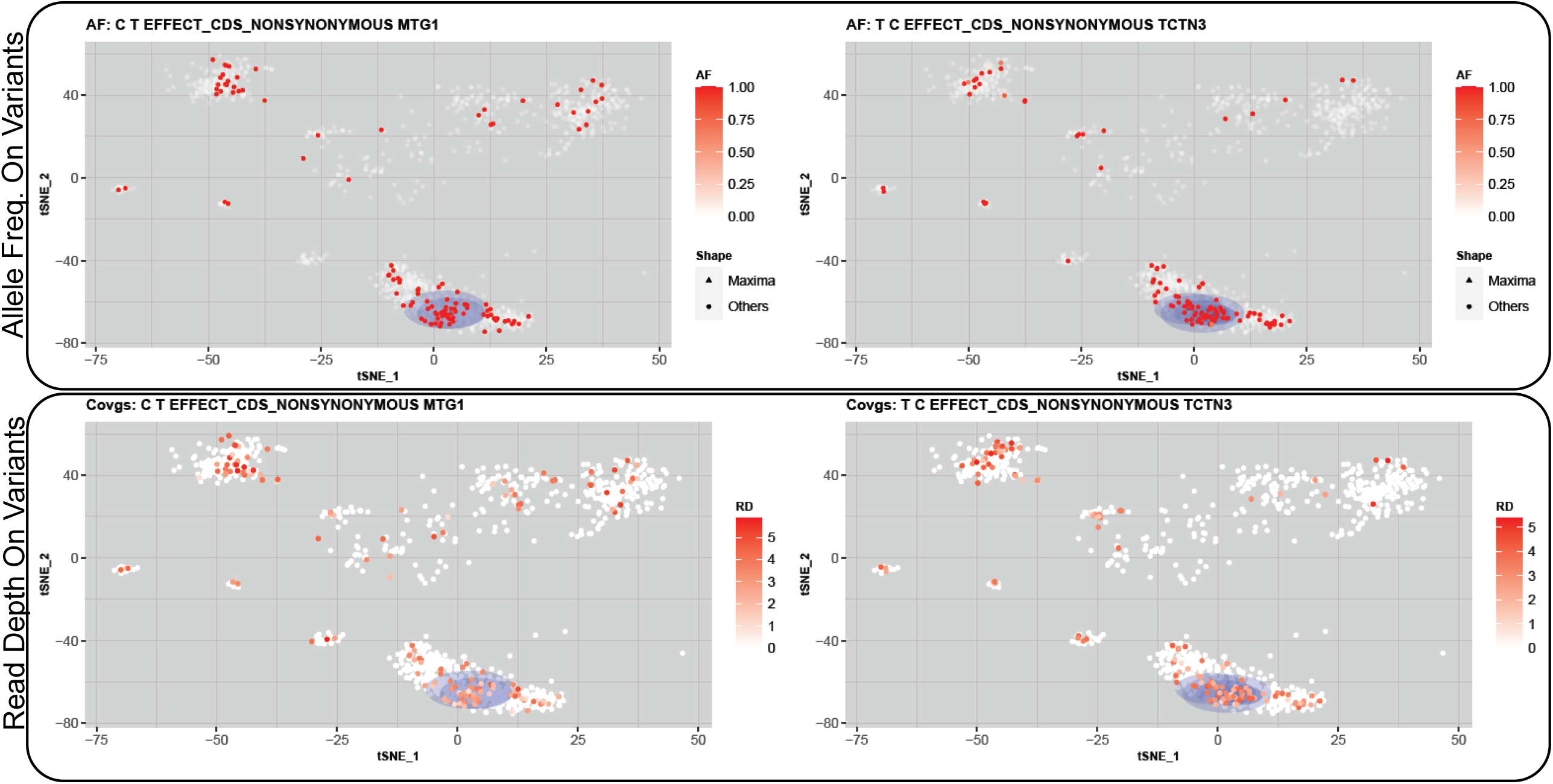
The analysis of small and large scale CNV clumps from Darmanis et al single cell RNA-seq dataset.

We next analyzed the SNVs and indels identified by XCVATR. One of the top deleterious variants that is detected by XCVATR is on TP53, which codes for a well-known DNA-repair protein (Figure 3c). This figure also shows the smoothed allele frequencies on all the cells. This mutation is also found in the COSMIC catalogue and is marked as deleterious. The smoothed signals show that there is a clear enrichment of the alternate allele frequencies among the malignant cells. One of the main aspects of clump detection is the read-depth at which each cell harbors the variants, which is shown in the bottom panel in Figure 3c. We also identified that several other genes, including TCTN3 and MTG1, which are highlighted in Figure 3c as forming significant clumps on the same set of cells. Interestingly, these genes are also mutated in some of the normal cells as it is seen on the tSNE embedding. These genes have been implicated with tumor biology in previous studies. These results highlight that clump detection can provide additional insight in analysis of tumor RNA-sequencing datasets.

### Bulk RNA-Sequencing from 160 Meningioma Samples

We finally used XCVATR to analyze the SNV and indels in bulk RNA-sequencing datasets. For this, we used an existing bulk RNA-sequencing dataset from a cohort of 160 meningioma patients^37^. We used XCVATR to detect and annotate variants (See Methods) and filter with respect to impact and population frequency (See Methods). Next, the identified variants are summarized to gene-level events. We generated the gene expression matrix and performed tSNE to generate the embedding of the data. We next ran XCVATR to identify the strong variant clumps. In order to evaluate the effect of tSNE parameters (namely the perplexity parameter, number of top variable genes, and minimum expression cutoff) that are used to detect the clumps, we ran tSNE on 180 different parameter combinations and ran XCVATR in each of the embeddings. We next analyzed the genes that are associated with the top scoring clumps (Fig 4). We saw that the top scoring clumps are associated with KLF4, AKT1, and TRAF7 gene mutations, which are most frequently reported by XCVATR among 180 different embedding parameters that were used. These three mutations are also reported as recurrent events in meningioma tumors^44^. Interestingly, the clumps associated with NF2 mutations (which are strong and most common drivers of meningioma) were scored lower in the XCVATR’s clump analysis (Fig 4). This result provides evidence that XCVATR can help uncover biologically relevant mutations in bulk RNA-seq samples.

**Fig. 4.**
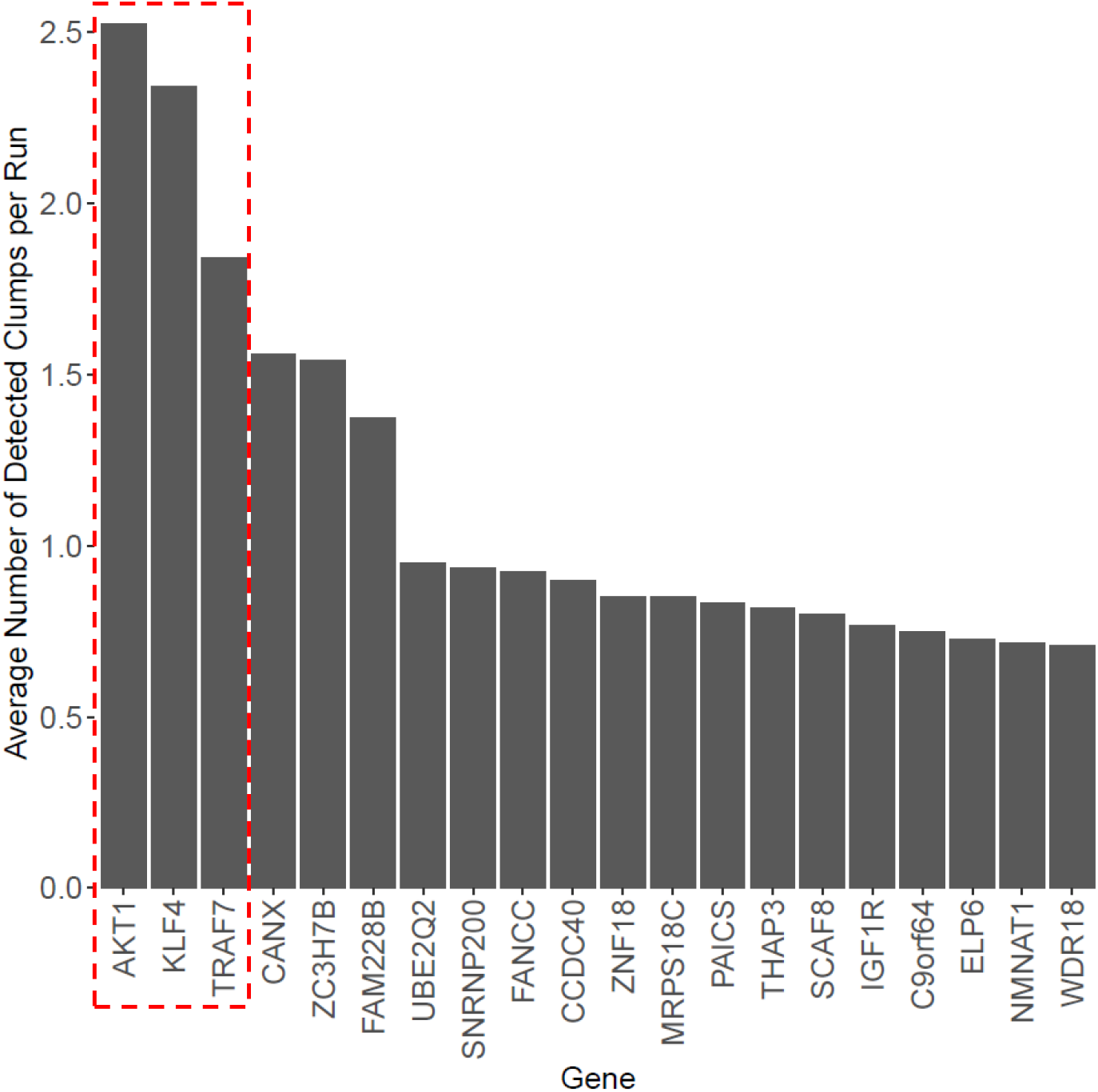
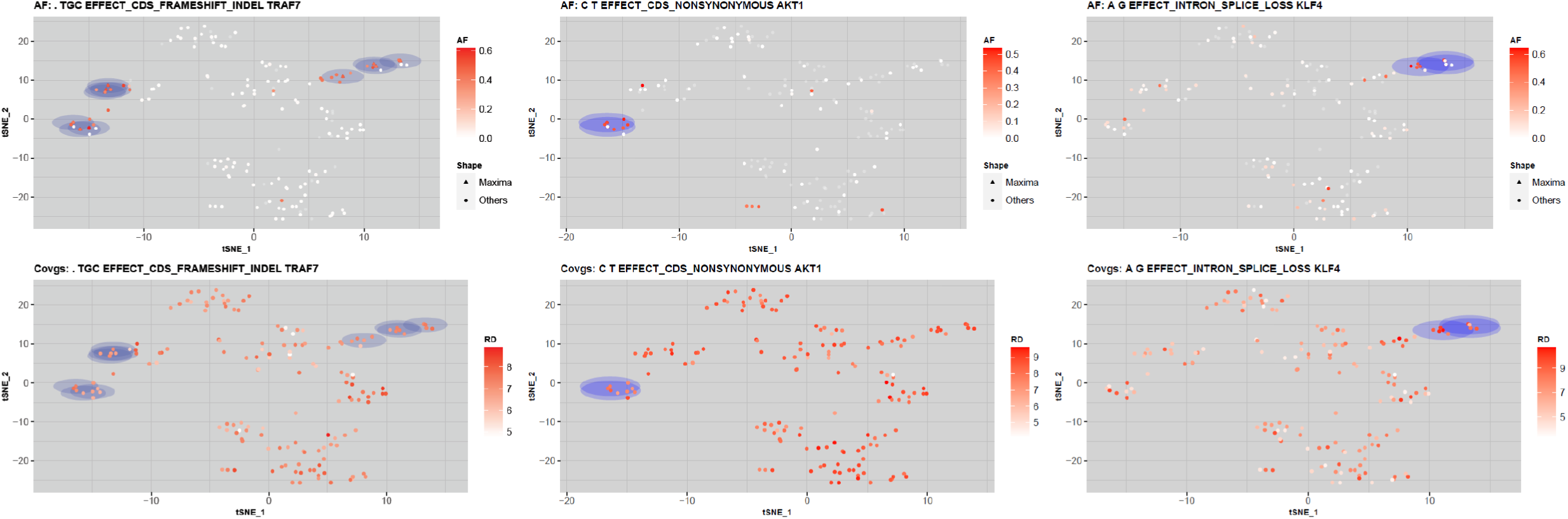
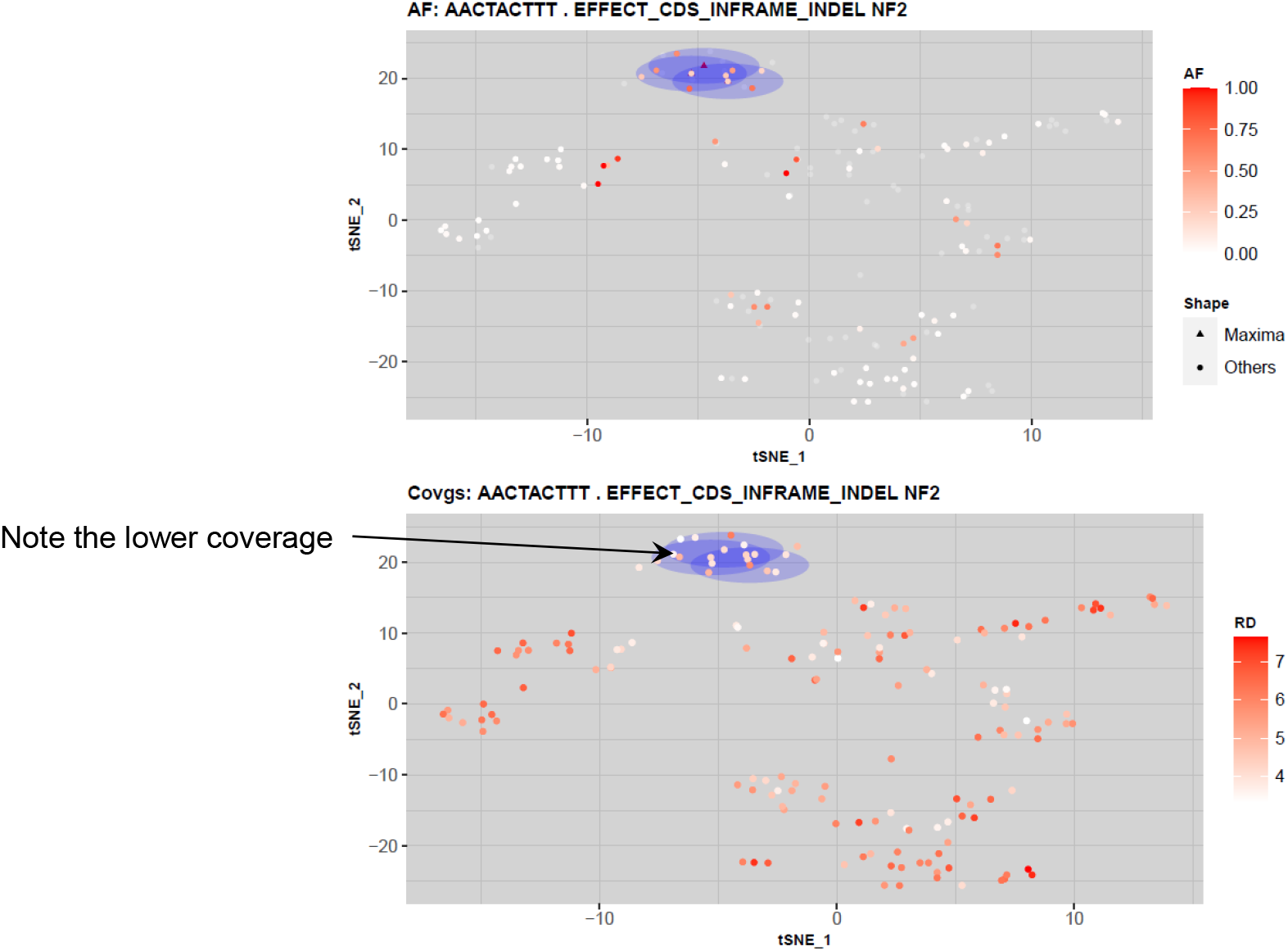
The analysis of the expressed variant clumps in 160 meningioma RNA-sequencing datasets.

## Discussion

We presented XCVATR, a method that analyzes the spatial enrichment of expressed variant alleles, i.e., local clumps in the cell and sample embeddings. XCVATR makes use of local spatial geometry of the embedding and multiscale analysis to provide a comprehensive workflow for detecting expressed variant clumps. These clumps can be used provide insight into driver genes and mutations by visualizing them on the embeddings, if possible. To provide as much control as possible, XCVATR integrates the variant detection, annotation, filtering, and clump detection in one package. This way, XCVATR does not explicitly have any requirements on variant calling methods such as GATK^45^. We hypothesize that this high level of control on variant calls is especially important since the variant calling can be made as relaxed as possible to provide a comprehensive set of variant calls, which can be stringently filtered at the clump detection stage. This way, rare somatic variants in cancer samples can be more inclusively analyzed compared to other existing pipelines that may miss rare variants with the default calibrated parameters^46^. One of the current limitations is that the concept of a clump needs more refinement and should be more discretely defined. In addition, the different embedding strategies should be surveyed to evaluate how embedding strategy and dimensions impact the clump identification.

## Methods

XCVATR algorithm’s methodology is summarized in Figure 1. XCVATR takes BAM formatted mapped read files. The bulk datasets that contain many BAM files, a BAM file per sample, can also be input to XCVATR. XCVATR makes use of samtools to process the BAM files and depends on a samtools installation.

### Quantification and Distance Matrix Generation

First, XCVATR quantifies reads on each gene for each cell (or sample). XCVATR makes extensive use of “CB:Z:” tag that is assigned by CellRanger software suite for assigning reads to different cells. This tag is also used internally by XCVATR to process bulk samples so that same implementation can handle single cell and bulk samples. The count matrix is used for generating the embedding of the cells in lower dimensions that will be used in the detection of variant clumps on the embeddings. The count matrix can also be used for building cell-to-cell distance matrix that can be used by XCVATR. Currently, XCVATR can provide tSNE and UMAP^47^ based embeddings of the cells into lower dimensions and XCVATR uses the distances from these embeddings. For single cell datasets, XCVATR uses SEURAT package to generate the tSNE/PCA/UMAP-based embeddings. For bulk samples, XCVATR utilizes “rtsne” function.

### Variant Detection and Annotation

Next, XCVATR performs detection and annotation of the genetic variants. This step can be optionally skipped in case there is a variant call set (i.e. VCF file) generated by other pipelines (GATK^28^, Mutect^29^). The users can provide these as input and skip the variant detection step.

### SNV Detection

XCVATR makes use of pileups to identify SNVs. To identify SNVs, the deduplicated (samtools) and mapping quality filtered reads (mapQ>30) from all cells are used to generate a pileup of the nucleotides at each position on the genome. Next, the candidates are filtered with respect to a minimum alternate AF cutoff. XCVATR also filters variants with respect to strand bias. For this, XCVATR build strand specific pileups and analyzes these pileups jointly to make sure the identified variants are not technical artefacts.

### Indel Detection

XCVATR performs scanning to identify the reads that support indels and clusters them to identify insertions and deletions. These are filtered with respect to mapQ and allele frequency.

### Variant Annotation

XCVATR takes variant annotation file (GTF or GFF) and annotates the variants with respect to their impact on the protein sequence. XCVATR maps the variants onto the transcripts that are specified on the GFF files. They are classified with respect to their location: CDS, splice, start/stop codons. For the mutations on the CDSs, the mutations are mapped on the coding sequences and coding impacts are evaluated: For SNVs, the variants are classified into synonymous, non-synonymous, splice altering, start/stop loss/gain. These are the most important impacts that we handle in XCVATR. For indels, the variants are classified into frameshift/in-frame (CDS overlapping length is multiple of 3 indicates in-frame), splice altering, start/stop loss.

### Allele Counting

For SNVs and small indels, XCVATR counts the number of mapping reads, for each read, that support the alternate and reference alleles. This is used to generate the estimated alternative allele frequency of each variant. XCVATR treats the alternate allele frequency of a variant These are then used as scores for a variant’s existence on each cell.

For copy number variants (CNVs), the variants are first separated into amplifications and deletions (Fig 1c). CNVs are different from small variants since they can cover large domains as long as the chromosomal arms. To analyze different length scales, XCVATR performs clumping analysis for large scale (chromosome arm length) and also at segment level scale. The large scale CNV analysis, there are 44 possible events for each deletion and amplification. For these, XCVATR first builds a binary count matrix that is analogous to the alternate allele frequency for the small variants. Each entry in this matrix indicates the existence of the CNV (row) in the corresponding cell (column). At the segment scale, each CNV is treated as a separate variant. However, since the CNVs identified in each cell has different coordinates, XCVATR first identifies the common amplification/deletion events by overlapping the CNVs from cells and identifying the minimal set of common variants (Fig 1). Next, these common variants are used to build a binary count matrix similar to the large-scale matrix. XCVATR analysis each of the common and disjoint events as a separate variant and performs variant clump analysis.

### Gene Level Summarization of SNVs/Indels

After the allele counting, XCVATR iterates over each cell and each gene and assigns the highest impacting mutations allele count to this gene: 

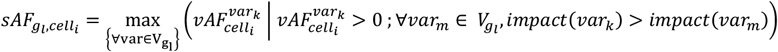

The gene-level summarization takes into account the positioning of the variants and removes some of the information. It is worth noting that summarization is an optional step as the clump detection can be performed at the variant level.

### Smoothing Scale Selection on the Embedding

XCVATR performs a multi-scale analysis of the distances to identify the variant clumps. Each scale defines a neighborhood around a cell in the embedding coordinates and is used to smooth the allele frequencies using a Gaussian filter that is centered on a cell and decreases as the cells get further from the center cell. The scales are tuned to the distance metric or the embedding coordinates. XCVATR performs a scale selection to tune the analysis to the selected cell-cell distance metric.

For each cell, XCVATR identifies *N*_*v*_ cells that are closest to the corresponding cell (i.e. neighbors). This defines the close neighborhood of each cell. XCVATR then scans the neighborhood size 

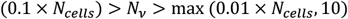

between 1% of the cells (or 10 cells, if lower) and 10% of the cells in the sample (*N*_*cells*_) and computes the radius of the neighborhood of each. For *N*_*v*_ cell neighborhood of a center cell, neighborhood radius is defined as the distance of the furthest cell to the current center cell. The minimum and maximum scales (*σ*_*min*_, *σ*_*max*_) are defined as the median of the neighborhood radii of all cells computed at the minimum and maximum neighborhood sizes *N*_*v*_ defined above (See Methods). The scales are used for smoothing the allele frequencies and identifying the variant clumps.

This computation can be performed efficiently since the distance matrix (unless it is provided) can be computed quickly from the embedding coordinates using fast matrix multiplications. Neighbor detection is performed by sorting the distances and selecting the closest *N*_*v*_ cell (or samples). After this step, only closest neighbors are processed by XCVATR.

### Variant Clump Candidate Selection

One of the challenges in clump detection is the large number of cells that needs to be analyzed in different scales. To decrease the search space, XCVATR performs a cell-centered analysis, where XCVATR does not aim at modeling the geometry of the embedding space but rather focuses on the cells, i.e., each detected clump is centered around a specific cell (or sample). We believe this is a reasonable expectation since the expected clumps are substantially larger than the cell-cell distances and therefore clump-detection should be accurate even when they are centered around cells. This way, XCVATR cuts the cost of modeling and searching the whole embedding space and focuses on cells.

Secondly, in visual evaluation of the variant allele frequency distributions on embeddings, the number of clumps were observed to be much smaller than the number of samples or cells. Motivated by this, we designed a candidate pre-selection that decreases the search space for the variant clumps. Given a smoothing scale *σ*_*a*_ at the scale *a*, XCVATR computes a smoothed AF value for each cell: 

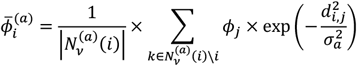

where *ϕ*_*j*_ denotes that alternate AF of the variant on the *j*^*th*^ cell (1> *ϕ*_*j*_ >0), *d*_*i,j*_ denotes the distance between *i*^*th*^ and *j*^*th*^ cells in the sample, 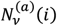 indicates the set of indices for the cells that are in the vicinity of the *i*^*th*^ cell for the scale *a*. From the above equation, the smoothed allele frequency of *i*^*th*^ cell, 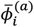, is higher when its neighborhood contains many cells with high allele frequencies. In addition, the smoothed AF depends on the scaling parameter *σ*_*a*_. Each scale is processed independently from other scales. The cells with high smoothed allele frequencies represent the potential variant clump centers in the embedding coordinates. XCVATR identifies the set of cells as candidates for which the cells in the neighborhood are strictly lower in terms of smoothed allele frequency (Figure 1): 

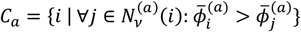

where *C*_*a*_ denotes the indices of cells that are clump centers. Above condition defines the candidate clump centers as the cells which have locally maximum smoothed allele frequencies when they are compared to the smoothed allele frequencies assigned to all their close neighbors.

### Specification of Position and Size of Clumps

Up to current point, we identified a clump by the cell at its center, which specifies the position of the clump in the embedding space. In addition to the center, it is also necessary to define the radius of the clump so that the size of the clump can be specified. XCVATR makes use of the scale parameter at which the clump is identified, i.e., *σ*_*a*_ at scale *a*. Thus, all the cells that are closer than *σ*_*a*_ to the center of a clump are assigned to this clump. Later on, we provide a method to detect the most enriched

### Variant Clump Evaluation by RD-aware permutation

For each of the cells in *C*_*a*_(*a*^*th*^ scale), the smoothed allele frequencies are compared to an empirical background. XCVATR utilizes a permutation test to assign significance to the each of the candidate clump centers. For this, XCVATR generates *K* permutations of the AF’s that are assigned to the cells and computes the smoothed AF for all the candidate. For each permutation, the smoothed AFs are computed for each candidate clump center. XCVATR then computes a z-score that is used to rank the clumps. 

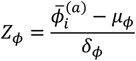

One of the important factors is to ensure that the AF provides new information and does not simply recapitulate the coordinates-based information in the embedding. One example for this is that some cells may exhibit a cell-specific marker that is not expressed in other cells at all. In this case, a germline variant will be expressed on these cells while other cells will have no expression of the variant. In such a scenario, a naïve approach would determine that this variant exhibits a clump on the cells where the gene is expressed. This would be an uninteresting clump that emerges based on the cell-type specificity of the gene. XCVATR aims at uncovering variant-specific clumps. To filter out these clumps, XCVATR sets a threshold *τ* on the total read depth at which the variant is expressed. XCVATR also reports the read depth z-score. This way, the clumps are evaluated with respect to the read-depth bias.

### Read-level and Cell-level Enrichment of the Alternate Allele Expression in the Clumps

In order to filter the clumps, XCVATR computes the significance of enrichment of expressed alternate alleles at the read level and at the cell (or sample) level in each clump. To compute the enrichment at the read level, XCVATR first computes the total number of alternate and reference reads in all cells. These are used to estimate a baseline (bulk) alternate AF. Next, for each clump, the total alternate allele supporting reads and total reads are computed. At scale *a*, and the *b*^*th*^ clump, these are used to compute the read-level modified binomial p-value: 

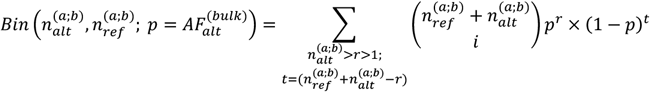

where 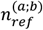 and 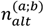 denote the number of reads that support alternate and reference alleles for the corresponding variant in the *b* clump that is identified in scale *a*: 

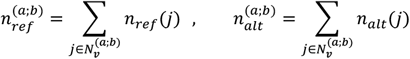

where 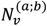 denotes the neighborhood of the center of the clump *b* at the scale *a*, and *n*_*ref*_(*j*) indicates the number of reference alleles in cell *j*. In above equation, the flip probability is chosen as 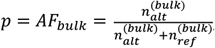, which represents the alternate allele frequency of the variant in the whole bulk sample. The binomial p-value estimates the enrichment of the alternate allele supporting reads in the clump *b* when compared to randomly assigning reads to all cells with probability *p* = *AF*_*bulk*_.

Next, XCVATR computes enrichment of alternate AF at cell level. At the scale *a*, XCVATR counts the cells in the clump *b* whose alternate allele frequencies are above *η*. Next, XCVATR counts the number of cells in the whole sample for which the alternate allele frequency is above *η*. These values are used to compute a significance of the enrichment of alternate alleles at cell level using Fisher’s exact test: 

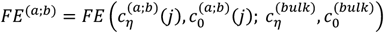

where 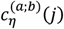 indicates the number of cells in clump *b* in scale *a* where allele frequency exceeds *η*: 

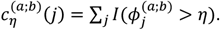

Similarly, 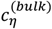 indicates the number of cells among all the cells (i.e., bulk) for which the allele frequency exceeds *η*. The read-level and cell-level enrichment estimates are used to filter out clumps that exhibit low levels of enrichment in comparison to the bulk sample at read and cell (or sample) level.

Finally, XCVATR computes the effective radius for each clump. For each clump, XCVATR iterates over the cells closest to the clump’s center cell. This way, the neighborhood around the clump’s center are analyzed as they expand in radius. For each neighborhood, the neighborhood where the cell-level enrichment is maximized (Fisher’s exact test p-value is minimized) is selected as the effective radius of the clump. After the clumps are identified, the clump centers and the scale at which they are identified, permutation z-scores, and alternate allele enrichment statistics, and effective radii are reported in the output.

### Visualization

XCVATR provides visualization of the clumps on the embedding coordinates for each variant. This enables the users to manually evaluate the variants. This can also be helpful to visualize the cell-type specifications and phenotypic properties in comparison to the clumps. The visualization utilities are implemented in R and directly make use of the data generated by XCVATR.

### CNV Calling by CaSpER

We have used CaSpER^9^ for detecting the copy number variants.

